# Membrane state diagrams make electrophysiological models simple

**DOI:** 10.1101/051839

**Authors:** Robert Law, Stephanie R. Jones

## Abstract

Ion channels are ubiquitous in living systems. Through interactions with membrane potential, ion channels both control metabolic events and mediate cell communication. Consequentially, membrane bioelectricity bears on fields ranging from cancer etiology to computational neuroscience. Conductance models have proven successful in quantitatively capturing these dynamics but are often considered difficult, with interpretation relegated to specialists. To facilitate research in membrane dynamics, especially in fields where roles for ion channels are just beginning to be quantified, we must make these models easy to understand.

Here, we show that the membrane differential equation central to conductance models can be understood using simple circular geometry. The membrane state diagrams we construct are compact, faithful representations of conductance model state, designed to look like circular “cells” with currents flowing in and out. Every feature of a membrane state diagram corresponds to a physiological variable, so that insight taken from a diagram can be translated back to the underlying model. The construction is elementary: we convert conductances to angles subtended on the circle and potentials to radii; currents are then areas of the enclosed annular sectors.

Our method clarifies a powerful but prohibitive modeling approach and has the potential for widespread use in both electrophysiological research and pedagogy. We illustrate how membrane state diagrams can augment traditional methods in the stability analysis of voltage equilibria and in depicting the Hodgkin-Huxley action potential, and we use the diagrams to infer the possibility of nontrivial fixed-voltage channel population dynamics by visual inspection rather than linear algebra.

## 1 Introduction

It is difficult to overstate the importance of ion channels in biology. They are nearly universally present at the threshold of living systems from the viral capsid [12, 48] to the mammalian cell membrane [36]. As mechanical [43], chemical [38], light [44], and temperature receptors [7], they constitute perhaps the largest part of cell sensoria. Ion channels form gap junctions [14], gate synaptic action [24, 11], and through direct ionic signaling [32] are essential mediators of communication among even bacterial populations. Voltage-gated ion channels are not only responsible for the neural action potential [18, 20] and muscular contraction [37, 13, 23, 2]: they also play crucial roles in the cell cycle and development, while abnormalities in ion channel expression are implicated in cancer [4, 31, 10]. Indeed, the mitotic mechanism itself is the target of a transduction pathway directly controlled by membrane potential [49]. Even the fertilization process [45, 46] depends on ion channel population and membrane voltage dynamics. Of all the topics in the life sciences, perhaps only genetics and proteomics can rival this functional span.

Conductance-based modeling is an extremely powerful technique for studying membrane dynamics used widely in neurophysiology (where it was originally developed in [18, 20, 19]), as well as computational neuroscience [15, 21] and cardiology [35]. This mathematical formalism has existed for over 60 years, and it is remarkable that models of ion channel dynamics are not nearly as commonplace as, for instance, genetic models. With notable exceptions [10], though, the functional importance of bioelectricity has traditionally been circumscribed to “excitable” tissue readily capable of generating action potentials, namely neurons and myocytes. A much larger variety of cells are now known to be excitable: action potentials occur in bacterial cells [25] and a wide variety of mammalian cell types (reviewed in [3]).

Moreover, even so-called “nonexcitable” tissues that do not typically generate action potentials have been shown to exhibit nontrivial dynamical phenomena like multiple stable membrane potentials [1]. Conductance modeling has recently been used to show that these “voltage memories” can be mediated by taxon-specific classes of ion channels [8, 26], and that when combined with gap junction models, these dynamics can generate rich spatial patterning of voltage in tissue [9]. We can expect that as this approach matures, interest in conductance models will only grow.

A major obstacle in applying these models is their mathematical overhead. At the center of the conductance model formalism is a differential equation (Equation 1) describing how the membrane voltage changes as currents cross the membrane. These currents are themselves governed by the evolving conductances of ion channels, which open and close according to their own dynamical rules (and are typically ascribed their own differential equations), as well as the reversal potential, which captures thermal and electrical contributions to ion flow. Biophysically, this approach is quite natural. However, differential equations are far from a standard tool in biology, and it has been our experience that even computational neuroscientists can find conductance models difficult to master.

Our goal here is to render conductance models simple enough that they may be used and understood with little mathematical background beyond a familiarity with circular geometry. To that end we present in Section 2 a visualization method called a membrane state diagram, which captures a great deal of quantitative information about ion channel populations and electrical variables, all while carrying the appearance and intuitive ease of a circular cell with currents flowing in and out. Our construction relies on an elementary correspondence between the sector area formula for an annulus (Equation 4) and the formula relating current, membrane voltage, reversal potential, and conductance (Equation 2). A membrane state diagram is a faithful geometric representation of conductance model state at any point in time: voltages are radii, angles are conductances, and areas are currents. Because of this correspondence, intuition gained by examining membrane state diagrams can be mapped precisely back to mathematical models.

The reader is urged to compare these diagrams to the standard textbook treatment of conductance models involving circuit diagrams (Figure 1a) and heuristic depictions of ion concentration gradients (exemplified by Figure 1b). To illustrate the utility of membrane state diagrams, we examine in Section 3.1 voltage perturbations about equilibria in a model with two channel populations, where the method was first applied [26] as traditional visualization methods proved insufficient for linking channel state to dynamics. In Section 3.2, we depict several phases of an action potential in the classical Hodgkin-Huxley squid giant axon model. Finally, in Section 3.3 we pictorially demonstrate the possibility of nontrivial dynamics (e.g. oscillations) in ionic currents occurring when voltage is held fixed, and then construct a model for generating those dynamics.

**Figure 1:**
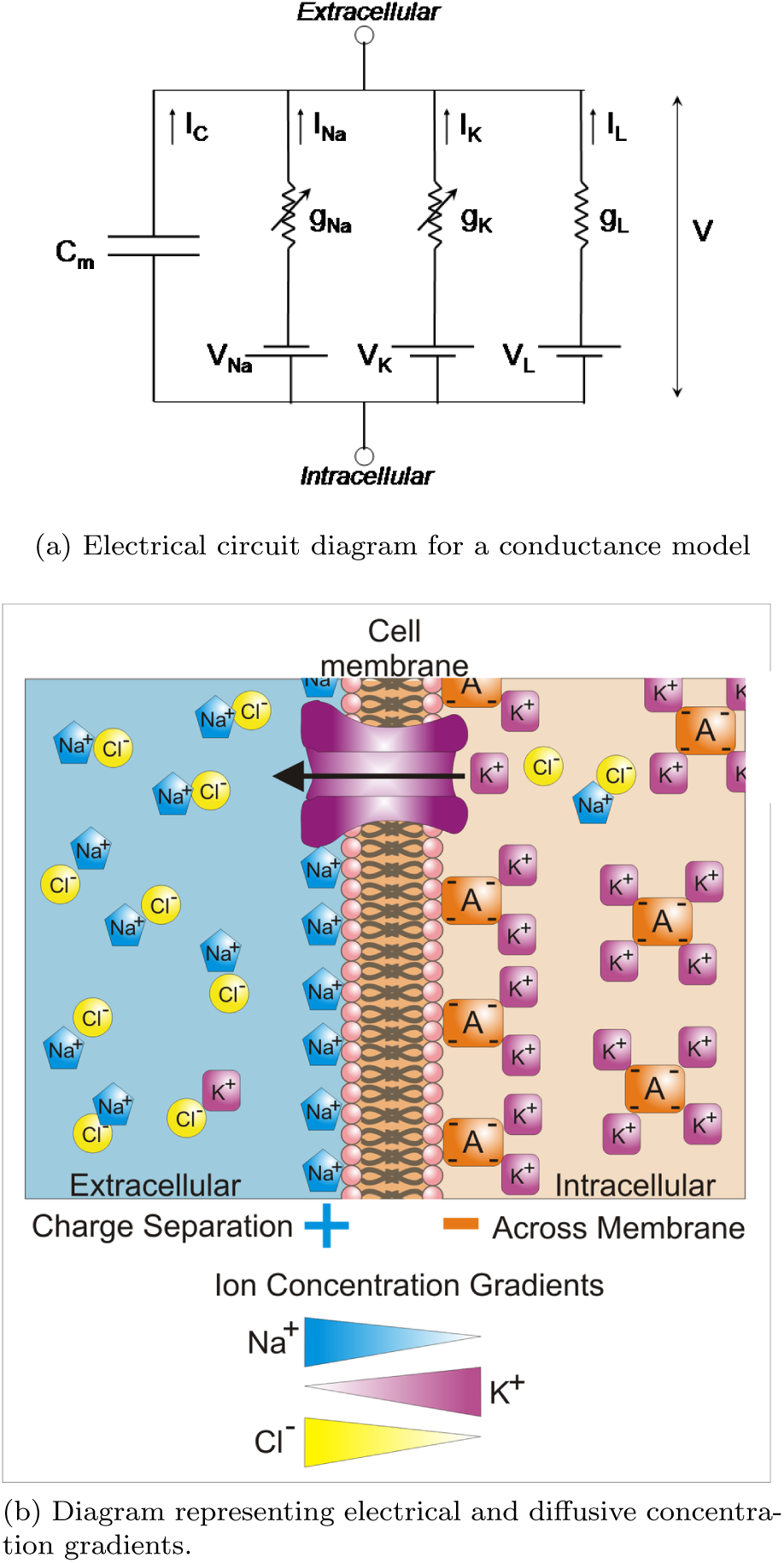
Traditional methods of depicting membrane dynamics do not allow for the accurate visualization of membrane state. (a) captures the model structure but does not depict state, while (b) partially captures state but cannot be mapped accurately to a model. (a) is from [39], (b) is from https://en.wikipedia.org/wiki/Membrane_potential#/media/File:Basis_of_Membrane_Potential2.png;author:Synaptidude; http://creativecommons.org/licenses/by/3.0/

## 2 Methods

Before introducing membrane state diagrams, we will begin with a brief treatment of conductance models. More detailed treatments may be found in [17, 6, 21]. The reader already familiar with these models may skip to Section 2.2, but should note our approach dispenses with finer points of channel activation and inactivation, simply referring to channels as “open” when conducting ions and “closed” when not conducting ions.

The diagrams can be applied *without reference to the underlying theory*. Once the membrane state diagram’s geometry is understood (angles representing channel populations, radii representing membrane and reversal potentials, areas representing currents), one can extrapolate dynamics knowing only one additional fact: that a net outward current leads to an increasing membrane voltage, while a net inward current leads to a decreasing membrane voltage.

Python and MATLAB functions for generating membrane state diagrams are documented and linked to in the Appendix, and videos depicting the dynamics of state variables can be found in the Supplementary Materials.

### 2.1 Conductance models

A conductance model treats a patch of membrane as a parallel electrical circuit (Figure 1a) consisting of a capacitor (the membrane surface, which collects free charge) and one or more transmembrane currents (ions crossing the membrane). With *Q* representing charge and *V* representing voltage, the capacitance is defined as *C = Q/V*, or the amount of charge separated by the membrane for each unit of voltage across it. If we assume the capacitance is constant, we can rearrange variables and differentiate with respect to time, yielding 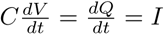. In other words the membrane voltage changes if and only if charges rearrange across the membrane, which itself forms a current.

Assuming this current crosses the membrane through ion channels (and is not instead bound, as occurs especially in the case of calcium; see Discussion), we use Kirchoff’s current law (*Σ I =* 0) to obtain the following ordinary differential equation:

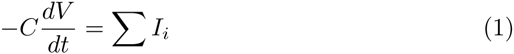

where *i* indexes a channel, and *I*_*i*_ is a transmembrane current through that channel. Equation 1 is simply a statement about charge conservation: an ion that crosses the membrane then collects on its surface, which changes the voltage according to the capacitance. When the current is zero, for instance, we see that *dV/dt* = 0; the voltage does not change because no net charge is accumulating on the membrane.

If we make a standard assumption that current flows according to Ohm’s law (see, however, [6, Ch. 2]), each current can then be written as

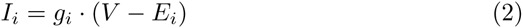

where *g*_*i*_ is the channel’s conductance and *E*_*i*_ its associated reversal potential. We will now examine the conductance and potential contributions to the current in turn.

#### 2.1.1 Conductances

A channel conductance *g*_*i*_ describes how permeable that channel population is to the ions it conducts. These conductances rise as channels open and fall as they close. The maximal conductance for channel type *i*, denoted *ḡ*_*i*_, grows linearly with the number of *i*-channels in the membrane, and *g*_*i*_ = *ḡ*_*i*_ corresponds to all these channels being open. Conversely, *g*_*i*_ = 0 when all *i*- channels are closed. As ion channels are governed by thermodynamic rules, a given channel will only have a *probability* of being open. The normalized variable *p*_*i*_ = *g*_*i*_/*ḡ*_*i*_ corresponds to the proportion of open channels to total channels of that type.

An important class of channels, the leak channels, do not normally close, so *g*_*leak*_ = *ḡ*_*leak*_. Generally, though, a wide range of factors may determine whether any given channel changes its state. Ligand-gated channels open or close when particular molecules (e.g. hormones) bind to them, while voltage-gated channels change their state due to the membrane voltage itself. Opsins, including the channelrhodopsin family [27, 5], open or close when exposed to light. The way a channel responds to a driving factor may be simple (e.g. “persistent” channels that open monotonically with increasing voltage) or it may be complex - for instance, sodium channels responsible for neural action potentials [20] first open and then close nearly as rapidly.

#### 2.1.2 Reversal potentials

For a given population of ion channels, the reversal or Nernst-Goldman potential [29, 16] is the membrane voltage at which the electrical and thermal diffusion gradients balance so that no net current crosses that channel population. The reversal potential condenses all temperature, concentration, and charge information into a single constant with units of voltage: without this simplification, it would be necessary to model ions, including their temperature-dependent rates of diffusion and their electrical interactions individually. Instead, the reversal potential can be treated as a bias or battery, driving current across the membrane unless it is exactly counterbalanced by the membrane potential. The net effect is this: *an ion channel’s current drives the membrane voltage toward its own reversal potential*. The magnitude of this drive at any point in time is assumed proportional to the difference *V - E:* the larger the imbalance of charge, the greater the electrical force and the larger the driven current.

To see this principle in action, we can examine a model with a single population of leak channels (we depict this using membrane state diagrams in Supplementary Video 1), with

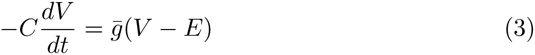

Whenever *V < E*, 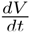 is positive, so voltage increases toward *E*. If *V > E*, the voltage drops, again toward *E*.

It is important to note that reversal potentials are associated to channels, not ions. Although we often assume that a channel conducts only one ion, many channels, for instance the HCN family [28], conduct more than one. A second important point is that unless ion concentrations are very low (as is the case in calcium in mammals), the amount of charge that crosses a membrane during even an action potential is typically so small that it has a negligible effect on the reversal (see e.g. [17]).

### 2.2 Equivalence of currents to annular sector areas

The key observation underlying our method is that the equation for an current across an ion channel (Equation 1) has the same form as the annular sector area formula:

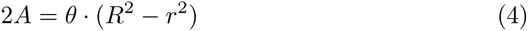

where *R* ≥ *r*. We will exploit this equivalence to generate a diagram in the form of a circular “cell” about which these sectors represent currents. This requires only a small amount of shoehorning in converting conductances to angles and voltages to radii.

#### 2.2.1 Representing conductances as angles

If one imagines a cell’s surface as a circle covered by ion channels, membrane conductances appear naturally as angles. A cell with 600 sodium and 400 leak channels, for instance, could be seen as having sodium channels on 3/5 of its surface and leak channels on the remaining 2/5. This geometrization does not map precisely to Equation 2, as a single open sodium channel is unlikely to have the same conductance as a single open leak channel. We may retain this intuition, though, and let each ion channel population occupy a fraction of the circle corresponding to that population’s contribution *ḡ*_*i*_ to the total possible membrane conductance *G* = Σ_*i*_ *ḡ*_*i*_.

We will associate the angles 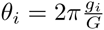 with the number of open channels and 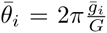 with the total number of channels of type *i* (Figure 2). For ease of notation, we suppress the normalization constant and refer to these angles as *g*_*i*_ and *ḡ*_*i*_ instead of *θ*_*i*_ and 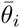. In a membrane state diagram, channel populations correspond to fractions of the “cell’s” surface, and currents will “flow through” a subangle representing the open subpopulation. This transformation “forgets” the total possible conductance G; we will remark further on this normalization in the discussion.

**Figure 2:**
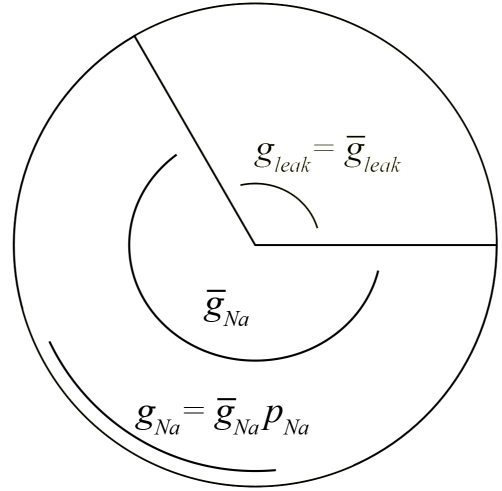
Angular representation of channel populations in a two-channel system with *ḡ*_*leak*_ = ½*ḡ*_*Na*_, *p*_*Na*_ = 0.4 and *p*_*leak*_ = 1. Viewing the circle as comprised of ion channels, the population of leak channels comprises one third of the total conductance and the population of leak channels comprises two thirds after normalization by 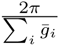 (see main text). The open sodium channels *ḡ* are represented as a subangle of the total sodium channels *ḡ*.

#### 2.2.2 Representing potentials as radii

Having associated angles to conductances, it remains to associate radii with potentials. The amount of current that flows through each population of open channels is proportional to the magnitude of the potential difference *V - E* (Equation 2), and the inward or outward direction of the current corresponds to its sign, where positive indicates current inflow and negative current outflow. In order to match Equation 4, it is the *square root* of a potential that we will associate to its radius in the diagram.

The reader may at first object to the possibility of negative arguments to the square root function, but note that we are free to “re-reference” V and E, shifting them to be nonnegative through *V ← V + V*_*shift*_ and *E ← E + V*_*shift*_. This does not affect the current, as first,

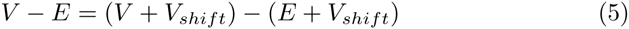

and second, although we do not unpack the conductances g_i_ here, these too must be invariant to voltage shift (in voltage-gated channels the half-voltage *V*_½_ plays a similar role to the reversal potential in Equation 5).

Because each current has *V* as a common variable, we set the radius of our p “cell” to 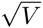. Each current then flows either inward or outward depending on whether *V* is larger or smaller than *E*_*i*_, so each radius 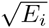 is either inside the cell (again, representing inward current flow) or outside the cell (representing outward current flow). Thus, in Equation 4, R will be the greater, and *r* the lesser, of 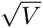 and 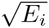. In this way, outward electrical drive is represented as pointing outward from the “cell’s” surface while inward drive points inward.

### 2.3 Membrane state diagrams

Each conductance angle and its associated voltage/reversal potential radii now bound an annular sector (shaded areas in Figure 3) oriented either outward (if *V* < E; negative current; Figure 3a) or inward (if *V* > E; positive current); Figure 3b). Each sector’s area is 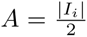, although we will again suppress the constant factor.

**Figure 3:**
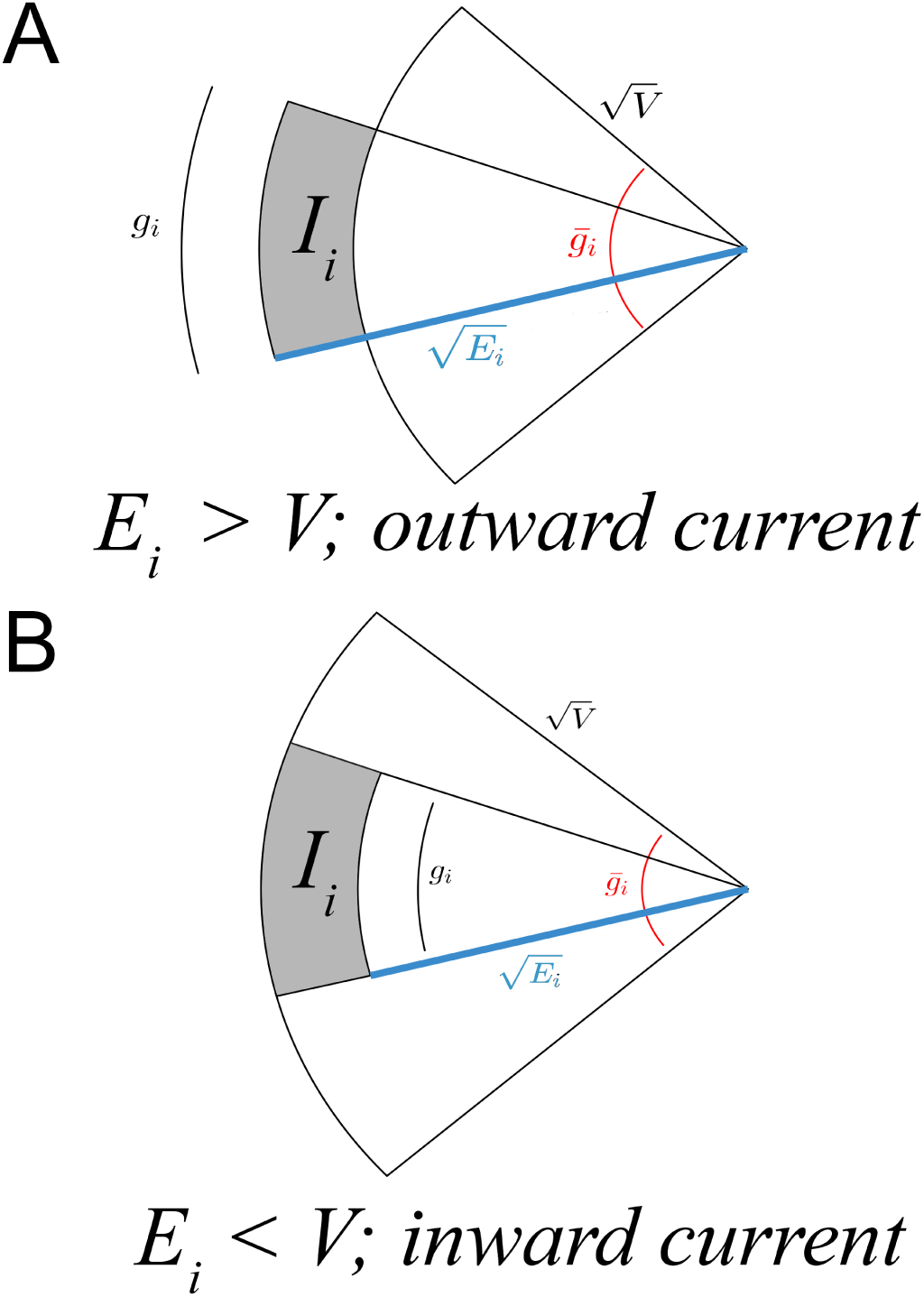
Schematic for the geometric representation of current across a channel population as an annular sector. As in Figure 2, each channel population indexed by *i* subtends an angle *ḡ*_*i*_, and the subpopulation of open channels spans the enclosed angle *g*_*i*_. A) An outward current *I*_*i*_ (grey shading) has magnitude proportional to the area of the annular sector with inner radius 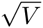, outer radius 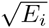 (blue), and angle *g*_*i*_. B) An inward current; same as (A) but as the membrane voltage is larger than the reversal potential, 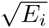 (blue) is the inner radius and 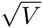 the outer radius. Factors of 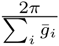 and ½ are suppressed in con ductance and current representations, respectively, and voltage variables have been shifted to be nonnegative; see main text. (A) is modified from [26]).

Assembling all these sectors into a disk, we now have a diagram (Figure 4) that simultaneously represents all membrane currents, ion channel conductances, reversal potentials, and membrane voltage while maintaining a strict correspondence with the mathematical model. In a membrane state diagram, currents are represented by areas of annular sectors that have radii corresponding to (square roots of) membrane voltage and reversal potential, while the sector angles are representative of channel populations.

**Figure 4:**
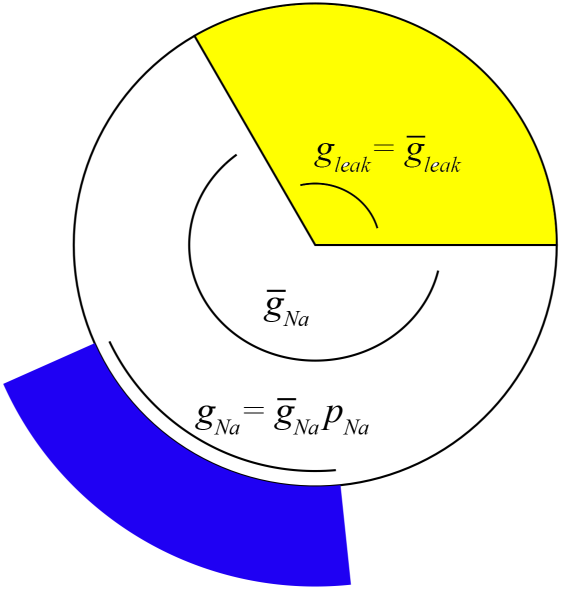
Membrane state diagram with same channel configuration as Figure 2. Here, *E*_*leak*_ is 0*mV* and *E*_*Na*_ is 100*mV*.

To infer membrane voltage dynamics from a membrane state diagram, note that a positive total transmembrane current (i.e. the right hand side of Equation 1) implies a negative 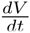. Thus, if the net current represented by the summed shaded areas is outward, the voltage increases and the ‘cell’ grows. Alternately, if it points inward, the voltage will decrease, and the ‘cell’ shrinks. At zero net current (inward and outward balance), the voltage does not change. For instance, a cell with only leak channels (Equation 3; see also Supplementary Video 1) is seen to always evolve toward its reversal. Membrane state diagrams also make it easy to see, for instance, that unless an external current is applied, the membrane potential is bounded between the highest and lowest reversals, and that synaptic inhibition by *GABA*_*A*_, which opens chloride channels, is excitatory whenever the membrane potential is below the chloride reversal.

Because membrane state diagrams are faithful representations of conductance models, they contain *only* the information already expressed in the membrane differential equation (Eq. 2). However, because the diagrammatic approach offers an integrated depiction of a large number of variables, it can clarify known model behaviors and lead to intuition about nontrivial phenomena that may not be easily inferred through examination of the equations.

We proceed with several more detailed examples to further illustrate the method.

## 3 Examples

### 3.1 Perturbations about equilibria in fast channel systems

Law and Levin [26] constructed and simulated conductance models of mammalian cells and amphibian embryos expressing a variety of ion channel combinations, first applying membrane state diagrams to graphically examine a stable voltage equilibrium in a fast voltage-gated sodium channel [40, 33] model with a leak current in mammals (cf. [21]). Fast channels have the property that the membrane voltage determines a unique channel conductance (i.e. the kinematics of the channel variables are rapid enough that they effectively reach their voltage-dependent equilibria instantly), so that at a given voltage, a system comprised only of fast channels has a uniquely determined membrane state diagram.

Here, we apply our method to visualize behavior around all three equilbria (two stable, one unstable) in this model. The conductance model for this system is:

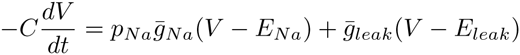

where *E*_*Na*_ = 60*mV* and *E*_*leak*_ = −67*mV*. The fraction of open channels is a sigmoidal function of voltage:

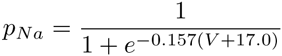

We overexpress the sodium channels relative to the leak channels, letting *ḡ*_*Na*_ = 10 · *ḡ*_*leak*_. The model’s phase portrait (expressing *dV/dt* vs. *V* and whose zero-crossings are voltage equilibria); top panel of Figure 5) shows stable equilibria near −67*mV* and 50*mV*, and an unstable equilibrium near −40*mV*.

**Figure 5:**
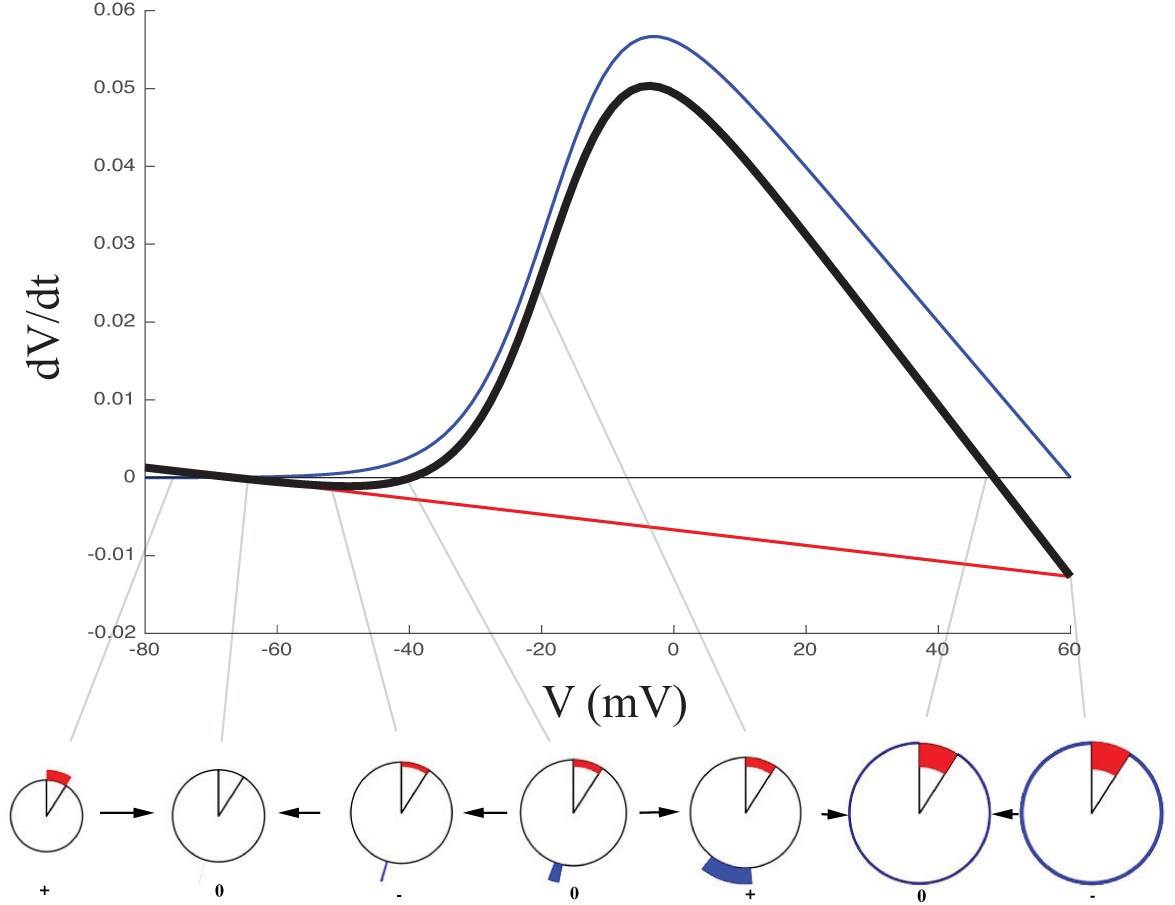
Top: Phase portrait for the mammalian fast sodium + leak model. Black is 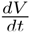 for the model; blue and red are the sodium and leak current contributions to 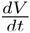, respectively. Bottom: membrane state diagrams at indicated voltages. Blue and red indicate the respective currents. The direction of voltage evolution is shown with black arrows, and the sign of *dV/dt* at each depicted voltage is indicated below each diagram (0 indicates a fixed point in voltage). Parts of this figure are modified from [26].

The phase portrait method (see e.g. [42, 21]) is indispensible for determining equilibria and allows for easy verification of stability (black curve in Figure 5; top panel). However, even after plotting the contributions of each ion channel individually (red and blue curves in Figure 5; top panel) it is difficult to see from a mechanistic perspective *why* these equilibria are stable. This issue is readily clarified by examining the membrane state diagrams at and near each of the three equilibria (Figure 5; bottom panel).

We can immediately see that the stable equilibrium at −67*mV* occurs nearly precisely at the leak reversal because almost all sodium channels are closed; this cell is then effectively equivalent to a cell with only a leak channel. In contrast, the other stable equilibrium at 50mV occurs near the sodium reversal because sodium channels are open *and* overexpressed relative to the leak channels (cf. [26]), making the equilibrium nearly equivalent to that of a sodium-only system. The unstable equilibrium near −40*mV* occurs where few sodium channels are open (small blue annular sector); a positive voltage perturbation opens more of these channels (blue annular sector increases in area), drawing the voltage toward the sodium reversal, while a negative perturbation closes sodium channels (blue sector decreases in area) and draws the voltage toward the leak reversal.

In this example, membrane state diagrams allow us to easily view features obscured in traditional analysis. The relative contributions of each channel population to the total current can be quickly read from the diagrams, and stability of each equilibrium can be both assessed graphically (stability means outward total current on the left and inward total current on the right) and linked to a particular ion channel configuration.

### 3.2 Hodgkin-Huxley system

We next use membrane state diagrams to visualize the channel dynamics underlying action potentials in a single-compartment Hodgkin-Huxley model axon [19], a conductance model obeying Equation 1 but with several nonlinear gating variables describing the kinetics of sodium and potassium channels. The sodium and potassium channels have conductances given by

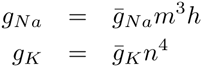

while the gating variables *m,n,h* ∋ *x* have dynamics governed by

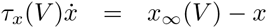

**τ*_x_* is a voltage-dependent time “constant” for channel gating kinematics and *x*_*∞*_ is the equilibrium state, where *τ*_*x*_ is Gaussian in *V* and *x*_*∞*_ follows a Boltz-mann function of voltage (see e.g. [21]; details may be found in the code linked in the Appendix).

Figure 6 shows the relative contribution of each ion channel at three phases of a simulated action potential. In this simulation, the action potential was initiated by setting *V*(0) to −35*mV*, above the threshold for action potential generation. Supplementary Video 2 depicts the membrane state over the entire course of an action potential. We see, for instance, that due to the rapid inactivation, at no point does the sodium channel population approach its maximal conductance.

**Figure 6:**
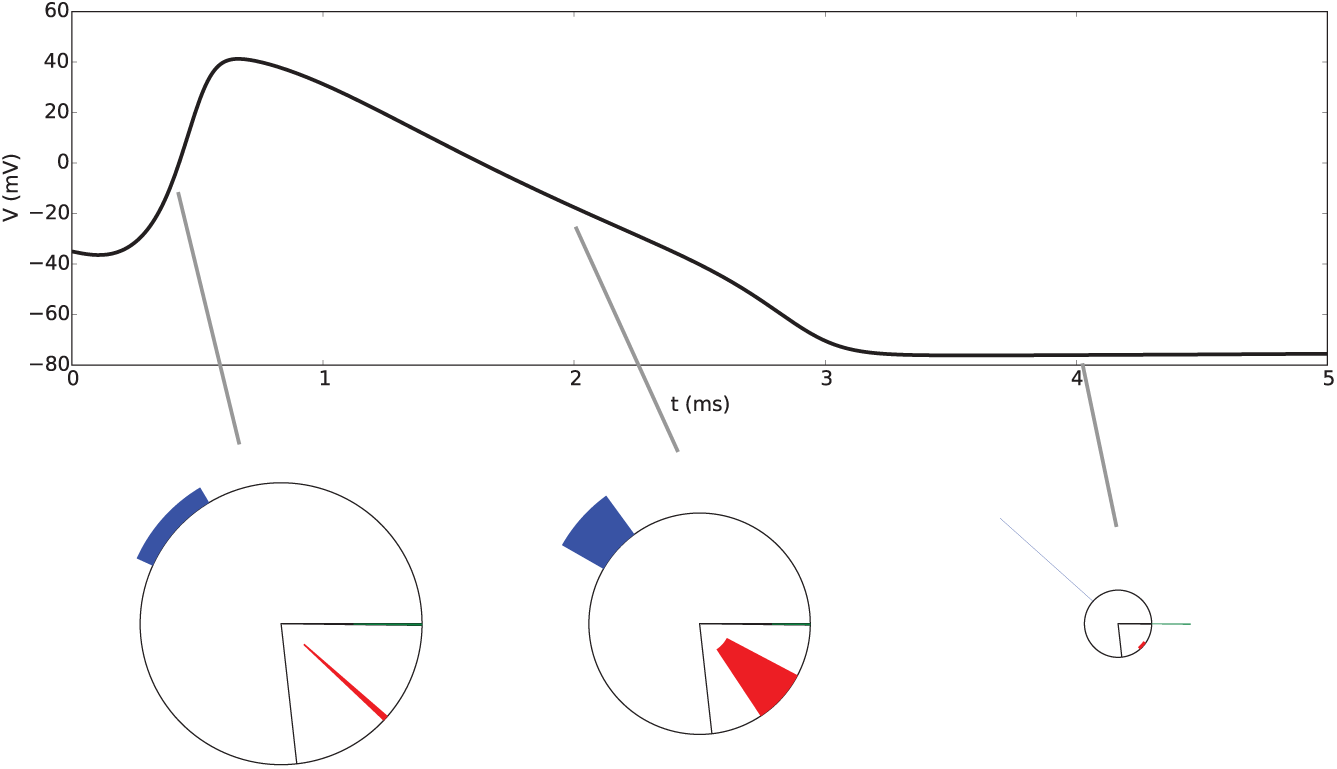
Top: Voltage evolution over time (not to be confused with the phase portrait in Figure 5) of an action potential in a Hodgkin-Huxley system. Bottom: Membrane state diagrams for this system at three points along the voltage trajectory: 0.5 ms (rising phase; left), 2.0 ms (falling phase; center), and 3.0 ms (recovery phase; right). Blue: sodium current. Red: potassium current. Green: leak current. Note that the sodium current is largest during the rising phase, and the potassium current in the falling phase. Due to its underexpression compared to the other ion channels, the leak current is negligible until nearly all the sodium and potassium channels are closed. A similar action potential is shown in Supplementary Video 2.

It is also seen clearly here that while the rapid rising phase of the action potential is caused by open sodium channels far from the sodium reversal, which establishes a large outward current, the falling phase is much slower due to a near balance of sodium and potassium currents. The leak current, expressed at dramatically lower levels than the other channels, plays very little role until the potassium channels close, and it finally overwhelms the potassium current in the recovery phase. To make a more general point, currents generated from small populations may be irresolvable visually in a membrane state diagram, but that this also reflects a negligible contribution to the overall current. Membrane state diagrams allow for these relative contributions to be assessed at a glance.

### 3.3 Channel dynamics under fixed voltage

Our third example shows how the formal correspondence between membrane state diagrams and conductance models may be leveraged to translate visual intuition into precise mathematical statements. Recall that when inward and outward currents are balanced (i.e. the net current is zero), the membrane potential does not change. A natural question to ask is, “Does fixed membrane voltage necessarily imply that all other system variables approach fixed points as well?”

In fact, this is not the case, and we can see easily (Figure 7) the possibility of ion channels “conspiring” to maintain zero net current, and therefore a fixed voltage, while undergoing complex dynamics of their own. For this to occur, it is merely required that changes in conductance (and/or in reversal potential, if concentrations are low) leading to an increase in current across one channel are balanced by changes leading to a decrease in current across other channels.

**Figure 7:**
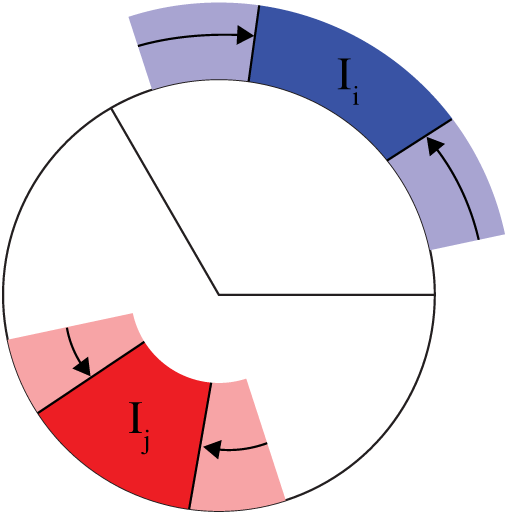
Membrane state diagram schematizing nontrivial channel population dynamics at a fixed voltage. Here, the outward current *I*_*i*_ is precisely balanced by an inward current *I*_*j*_ = -*I*_*i*_ at two time points (indicated by light and dark shading). Channels may open and close arbitrarily as long as they do so in proportion.

In other words, we have used the membrane state diagram to observe *entirely graphically* that the fixed-voltage condition is a linear constraint on (nonlinear) channel configuration and reversal potential dynamics:

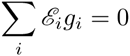

where *ℰ*_*i*_ = *V*^*^ - *E*_*i*_. This is of course easily verified by setting 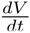 = 0 and *V* = *V*^*^ in Equation 1. This may occur in the two channel case where the voltage lies between the two reversal potentials and the channels open and close in phase.

Surprisingly, although it is quite easy to see that fixed-voltage oscillatory dynamics *may* exist, it is not entirely trivial to explicitly construct such a system. We do so here in the three channel case, although it remains to be seen whether these or similar dynamics occur in nature. Here, two channels will have outward currents and are reciprocally coupled to one another, forming a simple harmonic oscillator, while the third channel is an (inward) leak channel. We make the following assumptions. The outward populations have gating variables *x* with zero conductance when *x* = *x*^*^. We let *m*^*^ = *n*^*^ = ½. *g*_1_ = *ḡ*_1_(*n-n*^*^)^2^ and *g*_2_ = *ḡ*_2_ (*m-m*^*^)^2^. The shift is necessary to assure positivity of the gating variables, but an underlying mechanism may be conceptualized roughly as one where the channel is closed at 0 or 2 activated gates but open when 1 gate is activated, and where the number of activated gates tends to synchronize in the population.

We then have

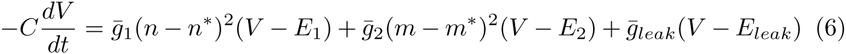

We assume *ḡ*_1_*ℰ*_1_ = *ḡℰ*_2_ = −4*ḡ*_*leak*_*ℰ*_*leak*_. At steady-state voltage, Equation 6 becomes:

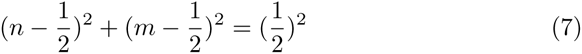

which is a circle of radius 1/2 centered at *n =* 1/2,*m =* 1/2. Assume reciprocal coupling of the form:

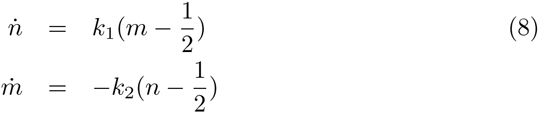

and let *k*_1_ = *k*_2_, concluding that if *m*(0),*n*(0) lie on the circle defined by Equation 7 and *ṁ*(0) and *ṅ*(0) lie tangent to it, the channel populations will oscillate at fixed voltage *V*^*^. While these oscillations are not stable to perturbations, a stable version can be constructed by using the Hopf oscillator equations (see e.g [34]) in place of System 8.

This example has shown that one can use visual inferences from membrane state diagrams to suggest processes that are not immediately evident in the conductance model formalism. These processes can then themselves be mapped back to equations, allowing for visual intuition to guide model development.

## 4 Discussion

### 4.1 Summary

Membrane state diagrams form a snapshot of ion channel configurations and currents at fixed time, allowing the viewer to simultaneously examine the contribution of ion channel population states and electrochemical potentials in generating membrane currents. We have applied this method to visualize channel dynamics during voltage perturbations near equilibria in fast channel systems, to detail the Hodgkin-Huxley action potential, and to make inferences about oscillating channel dynamics at fixed voltage.

### 4.2 Comparison with other approaches

While our method is not intended as a substitute for standard dynamical systems methods like phase portraits, it does augment these methods well when geometric intuition must be tied back to membrane state (e.g. Section 3.1). Membrane state diagrams were designed, rather, as a substitute for two textbook approaches: “cartoon” diagrams such as Figure 1b, which are suitably “biological” but cannot be used to make dynamical inferences, and circuit diagrams (Figure 1a), which schematically represent the biophysics but do not lend themselves to visual intuition.

When compared to bar-graph, tabular, or time series slice representations of current, membrane state diagrams may suffer from the “pie chart problem” [47]: it is typically more difficult to compare the areas of two pie chart sectors than to compare the same information displayed in a bar graph or table. None of these alternative methods, though, can simultaneously quantify conductance, distance from reversal potential, and current as well as the *relationships* among these quantities. Furthermore, to infer whether cell voltage is increasing or decreasing, the viewer must judge whether the *total* inward or outward currents are represented by larger areas. Spence and Lewandowsky [41] have found that when sums must be compared, pie charts afford *more* accurate comparisons than bar graphs, although it is not known whether this result would carry from circular sectors to annular sectors. In cases where it is difficult to judge whether the inward or outward current is greater, we recommend adjoining a bar graph representing the *summed* inward and outward currents next to a membrane state diagram.

### 4.3 Limitations and extensions

Due to the faithful correspondence of variables, the primary limitations of membrane state diagrams are limitations of conductance models themselves. One notable complication arising in these models is their treatment of calcium in typical mammalian intracellular concentrations^1^. Nonlinearities arising from calcium depletion and buffering can be mollified by directly modeling ion concentrations and then treating the calcium reversal as a variable. Similarly, voltage fixed points a la Section 3.1 may be “stable” only insomuch as the reversal potentials can be treated as constant. However, a 1:1 sodium/chloride leak exchange is clearly shown in the rightmost equilibrium in Figure 5, and at long timescales, these too must be treated by modeling concentrations directly. Membrane state diagrams do not keep track of these concentration variables, but may aid in visualizing their effect on reversal potentials.

However, because the diagrams are quite minimal and require no labeling intrinsically, they can be augmented to include additional information. As shown in Figures 5 and 7, arrows can indicate the direction of conductance changes or make the direction of voltage evolution more evident. When depicting a fast channel as in Section 3.1, the function *g*_*i*_(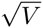) can be plotted radially as a guide, and finer channel properties (e.g. inactivation and deinactivation) of subpopulations could be indicated through, for instance, shading of the circle.

Any current modeled using Equation 2, for instance, a synaptic current, can be included in a membrane state diagram, but injected currents may also be depicted by recasting them as a conductance *g*_*I*_ and reversal *E*_*I.*_ However, the same injected current applied at two different voltages will not correspond to the same *E*_*I.*_ (and/or *g*_*I.*_)! Care must also be taken in setting *V*_*shift*_ to ensure that inward applied currents do not “overflow” the diagram. Alternately, applied inward currents might be assigned to the center of the circle. Again, this depends on a choice of *V*_*shift*_ that avoids overlapping currents. Although designed for biological contexts, membrane state diagrams can also be applied to represent current flow and voltage evolution in any parallel RC circuit with variable resistors, provided that the resistance of each element has a known lower bound (otherwise *ḡ*_*i*_ may be infinite).

### 4.4 Diagramming multiple cells and compartments

As constructed, membrane state diagrams describe “point cells”, single-compartment models with an isopotential surface and no spatial variation in channel expression [39]. They can, however, be used in a network context. The method may be particularly suited to depicting currents cylindrically for representing models that include axonal or dendritic processes (for instance, [22]), provided that the axial conductances are included in each segment.

When representing multiple cells or compartments, though, recall that the normalization procedure “forgets” a scale factor *C/G*. This means that two cells with identical membrane state diagrams need not actually have identical dynamics; however, their respective *dV/dt* will vary only by this scale factor. Given that cells appear to homeostatically regulate to maintain relative, but not absolute *ḡ*_*i*_ (see e.g. [30]), it may be the case that this scale factor is largely irrelevant for computation.

### 4.5 Conclusions

We expect that membrane state diagrams should prove particularly useful as a pedagogical tool. Once a student understands that an individual channel current tends to draw the membrane voltage toward its own reversal potential, the diagrams can be introduced to show conductance models in their full complexity with very little additional conceptual overhead. Membrane state diagrams, though, can facilitate insight into conductance models even among experts, and we anticipate that this technique will prove widely useful in the study of cellular physiology where it can complement or supplant traditional approaches to visualizing membrane biophysics.

## Acknowledgements

RL would like to thank Michael Levin for initiating the project that led to this work and for many helpful discussions, Erik Roberts for alerting us to the “pie chart problem”, and both Can Tan and the students of APMA2821v for reading an early version of the manuscript. Both authors thank the Jones lab for other valuable feedback. This work was supported by the National Institutes of Health (NIH MH106174).

## Appendix

The visualization method has been implemented in MATLAB and in Python, and it may be downloaded from https://bitbucket.org/nosimpler/msdiagram/. It consists of one main function:

**msdiagram** takes vector arguments *gbar, p, E*, scalar arguments *V* and *offset*, and in the Python case a matplotlib axis *ax*, returning that axis with a plot geometrically depicting the currents. Here, *gbar* is a vector of channel maximal conductances, *p* is a vector of channel open probabilities (for example, for the sodium channel in the Hodgkin-Huxley model, *p* = *m*^3^*h*), *E* is a vector of channel reversal potentials, *V* is the membrane voltage, and *offset* provides an additional shift in the voltage for ease of visualization (*V*_*shift*_ is computed such that *min*(*V, E*) = 0, and *offset* replaces 0 with a positive value). Code for generating movies depicting the Hodgkin-Huxley action potential may be found in the MATLAB version.

## Footnotes

1 This problem should not arise in e.g. amphibian embyros as the intracellular calcium concentration is quite high (see for instance [26]).

